# Tissue-specific and tissue-agnostic effects of genome sequence variation modulating blood pressure

**DOI:** 10.1101/2022.04.19.488795

**Authors:** Dongwon Lee, Seong Kyu Han, Or Yaacov, Hanna Berk-Rauch, Prabhu Mathiyalagan, Santhi K. Ganesh, Aravinda Chakravarti

**Affiliations:** Department of Pediatrics, Division of Nephrology, Boston Children’s Hospital, Boston & Harvard Medical School, Boston, MA, USA; Center for Human Genetics and Genomics, New York University Grossman School of Medicine, New York, NY, USA; Department of Internal Medicine & Department of Human Genetics, University of Michigan, Ann Arbor, MI, USA

**Author notes:** Send all correspondence to:* Aravinda Chakravarti, Ph.D., Center for Human Genetics and Genomics, New York University School of Medicine, 435 East 30^th^ Street, SM Room 802/3, New York, NY 10016, USA, (T), (E), Dongwon Lee, Ph.D. Division of Nephrology, The Manton Center for Orphan Disease Research Boston Children’s Hospital & Harvard Medical School 300 Longwood Ave, Enders 505, Boston, MA 02115, USA (T), (E). These authors contributed equally (Dongwon Lee, Seong Kyu Han).

## Abstract

Genome-wide association studies (GWAS) have mapped thousands of variants for numerous polygenic traits and diseases. However, with some exceptions, mechanistic understanding of which precise variants affect which genes in which tissues to modulate trait variation is still lacking. To this end, we introduce a novel class of genomic analyses applicable to any complex trait using GWAS together with gene expression and chromatin accessibility data from multiple tissues. Here we identify the transcription factors (TFs) and regulatory variants within active enhancers regulating specific genes in individual tissues to explain trait heritability of blood pressure (BP), a classical polygenic trait. We show that kidney-, adrenal-, heart-, and arterial-specific regulatory variants contribute to 2.5%, 5.3%, 7.7%, and 11.8% of variant heritability, respectively. Collectively, ∼500,000 predicted regulatory variants across these four tissues explain 33.4% of variant heritability. We demonstrate that these variants are enriched in enhancers binding specific TFs in each tissue. Our findings suggest that gene regulatory networks perturbed by common regulatory variants in a tissue relevant to a phenotype are the primary source of interindividual variation of BP. These studies provide an approach to scan each human tissue for its physiological contribution to a trait.

## INTRODUCTION

Genome wide association studies (GWAS) of numerous polygenic traits and diseases have been highly successful in mapping hundreds of thousands of common sequence variants clustered at thousands of genomic loci^1–3^. These genetic analyses have confirmed Fisher’s infinitesimal model of multifactorial inheritance^4^, namely, that complex traits and diseases arise from the additive actions of small genetic effects at hundreds to thousands of genes distributed across the genome^5^. The success in answering this genetic puzzle stands in stark contrast to our continuing inability to identify the mechanistic basis for this functional genetic architecture for the majority of polygenic traits. Most GWAS variants reside in the noncoding genome and are believed to perturb the expression of many genes by modulating the effects of transcriptional enhancers, likely in a tissue- or cell type-specific manner^6^. However, in order to understand the functional basis of a given trait or disease we need to know the identities of four components for each GWAS locus: the *transcription factors* (TFs) involved, the *cis regulatory elements* (CREs or enhancers) that bind these TFs, the *causal sequence variants* within CREs that alter enhancer activity and the *gene* whose expression is thereby altered^7^. Genetic variation in any or all components of this quartet is the fundamental genetic unit of polygenic traits. This problem has been difficult to solve, except for selected genes and loci^8,9^, and here we provide a general genomic and computational framework for doing so. When successful, these transcriptional components will allow experimental tests of the veracity of a trait’s uncovered *functional architecture* and teach us how a specific set of genes modulate a trait or disease.

Central to this task is the identification of the tissues where such genetic variation is active. We use the genetic principle that only sequence variants in CREs active in a specific tissue can mediate trait effects by perturbing their target genes’ expression in that tissue, irrespective of whether the target gene is expressed elsewhere. Thus, CRE activity in a given tissue is the most direct way to separate putative causal variants from those in linkage disequilibrium (LD) with causal variants. It is of course possible that a CRE sequence variant is merely in LD with the causal variant residing in another tissue’s CRE. Distinguishing these effects require us to predict the functional consequences of all CRE-resident variants, analogous to methods which distinguish pathogenic from benign coding variants. Advancing such analyses require comprehensive and high-quality tissue-resolved CRE and gene expression epigenomic maps which functionally constrains the genomic space where sequence variation relevant to a trait can reside. The availability of such maps will then allow scanning all human tissues for their relative contribution to a trait’s inter-individual variation, and, in principle, can be extended to all cell types within contributing tissues. The analyses we propose here are central to understanding biological mechanism because they connect genome sequence variation to cell and tissue effects of a phenotype.

Identifying functional variants in CREs is another major challenge because only a small proportion of variants within CREs affect its activities^10^. However, the advent of computational methods for functional CRE variant predictions now provides a practical means to systematically identify such variants genome-wide^11–14^. Over the past decade, several groups, including us, have pioneered these approaches and demonstrated that functional CRE variants can be predicted from DNA sequences.^15–17^ To this end, a two-step approach is typically employed. First, sequence-based models for CRE prediction are trained on experimentally detected CREs using machine-learning methods. Second, this model is used to estimate the effect of any variant within a CRE by calculating the difference in model-predicted CRE activities between the reference and variant alleles. This method can score any genetic variant as long as its sequence information is available regardless of its type, allele frequency and haplotype structure. This sequence-resolved epigenomic map can thus identify variants in CREs shared across tissues but affecting gene expression by perturbing the corresponding CREs in a tissue-specific manner. Thus, high-quality tissue-resolved CRE maps, also used as training data for these sequence-based predictive models, are an essential component of our analyses.

In this study, we propose genomic and epigenomic analyses of any complex trait and apply these methods to blood pressure (BP) variation, a classical polygenic trait. Genetic dissection of blood pressure has remained challenging because of the influence of numerous environmental and genetic factors, the mechanistic involvement of multiple tissues, such as kidney, adrenal, heart, and vasculature, among others, and multiple levels of physiological control, be they instantaneous, diurnal, or long term^18^. BP has been a major target of human adaptation in our evolution^19,20^ and is a significant physiological trait that underlies numerous cardiovascular diseases and their complications^21^. It is precisely because of the multiplicity of mechanisms that genetic analysis is critical because it can implicate genes, pathways and tissues, and their relative roles, in an unbiased manner. Genetic analyses of BP syndromes have clearly revealed the major role of salt-water homeostasis in the distal nephrons of the kidney as being critical^22^; however, BP epidemiology suggests that this is an incomplete picture, and many more tissues are important. The methods and studies we describe here, now allow us to answer these and many other broader questions regarding BP genetics and essential hypertension, one of the most frequent of human disorders.

## RESULTS

Our primary objective is to elucidate the functional genetic architecture of systolic (SBP) and diastolic blood pressure (DBP) contributing to interindividual BP trait variation in a tissue-resolved manner (**Figure 1**). We focus here on four major tissues relevant to BP^23^, the adrenal gland, artery, heart, and kidney, and first build high-quality epigenomic maps for gene expression, CREs, and regulatory variants (see **Methods**). Although many such relevant datasets are publicly available, they require uniform reanalysis because our tissue-resolved analysis is comparative and requires the use of identical criteria across datasets, and, our analysis relies on sequence-resolved data^11,24,25^.

**Figure 1:**
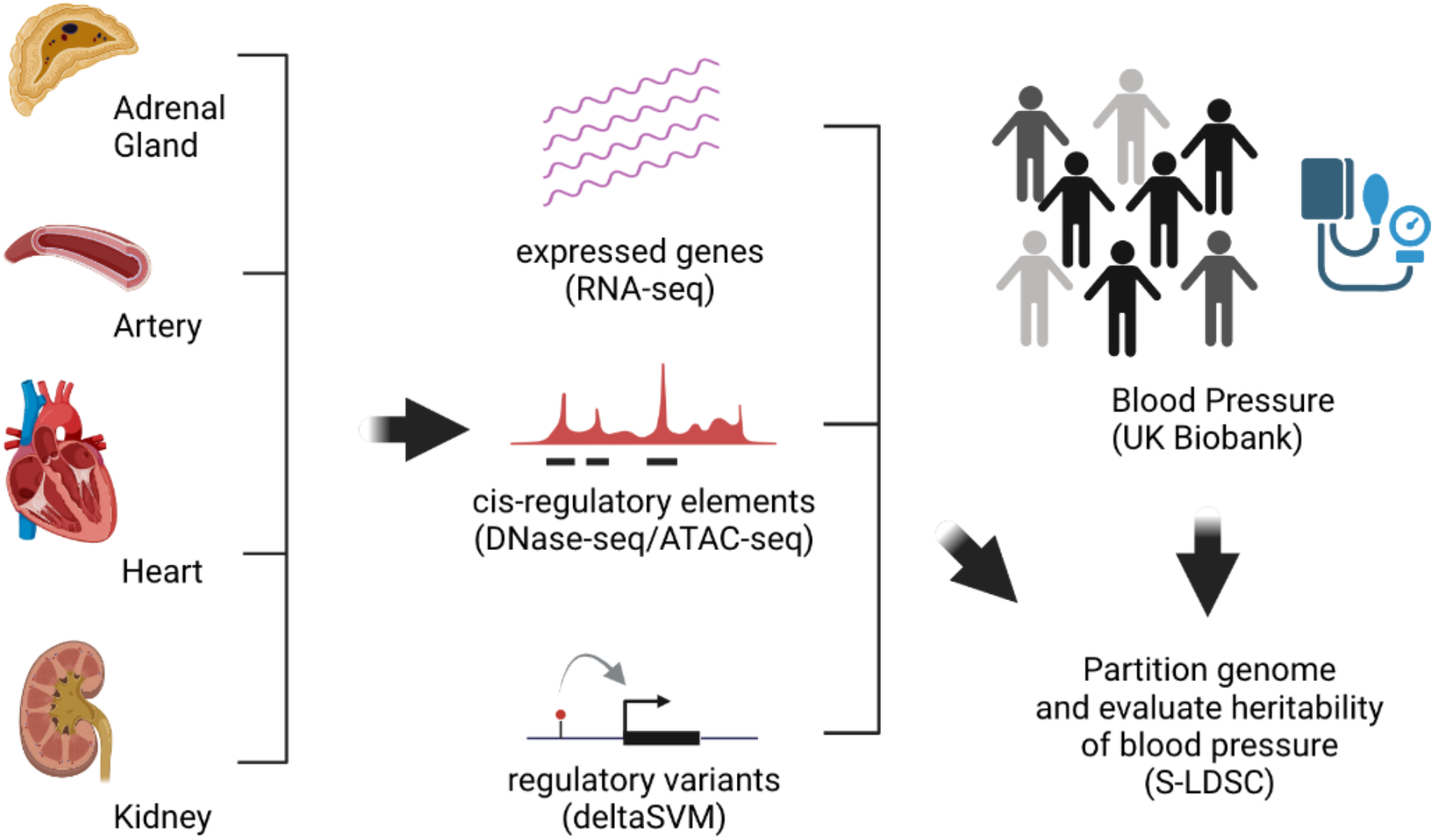
Study design. We focused on four major tissues relevant to blood pressure regulation and analyzed their relative contribution to systolic and diastolic blood pressure (BP) using genetic variants affecting gene expression of these tissues.

For CRE maps, we reanalysed existing chromatin accessibility data to construct comprehensive CRE maps. We first analyzed DNase-seq data for three adult tissues that were publicly available. Using our optimized peak calling methods for open chromatin^26^, we uniformly processed existing DNase-seq from the ENCODE project^27^ and compared our CRE peak calls to HOTSPOT2 peaks publicly available from ENCODE. We performed extensive sequence-based quality control (gkmQC) analysis, which we recently developed for assessing the quality of epigenomic data^28^, to show that our optimized methods identified higher-quality peaks (higher peak predictability) than ENCODE processing methods (**Figure S1**). Consistent with this comparison, the heritability of BP, estimated using the stratified LD score regression method (S-LDSC), is higher in our identified peaks than HOTSPOT2 peaks (**Figure S2**). For adult kidney tissues, we generated multiple ATAC-seq libraries as kidney open chromatin data were not publicly available (**Methods**). Along with our kidney ATAC-seq data, we uniformly processed additional ENCODE ATAC-seq data for other tissues as well and confirmed that several of them, including our kidney ATAC-seq, exhibited high quality and were comparable to DNase-seq data (**Figure S3**). Once again, heritability analysis demonstrated that high-quality ATAC-seq samples determined by gkmQC explains greater BP heritability than low-quality ones and were comparable to high-quality DNase-seq data (**Figure S4**).

Each sample for a tissue captures slightly different enhancer regions due to variation in technical (e.g., assay used) and biological (e.g., variation in CRE intensity) factors.^26^ Thus, it is necessary to construct comprehensive maps from all available chromatin accessibility data of specified quality for each tissue and analyzed in the same manner. Starting with the highest quality sample identified by gkmQC, we defined peaks in this sample as “the best peak set.” We identified unique peaks not overlapping these for each of the remaining samples from the same tissue and chose the next best sample whose unique peaks explained the largest additional BP heritability, from a partitioning heritability analysis (S-LDSC),^29^ and augmented the existing best peak set with these peaks. We repeated these steps until a sample failed to significantly increase explained trait heritability. This process showed that the two highest quality samples from different chromatin accessibility assays are often sufficient to explain the greatest BP heritability (**Figure S5**). We note that differences in the number of open chromatin regions across tissues could produce a potential bias in comparative heritability analyses. To reduce this effect, we selected the top 100,000 regions from each tissue for further analysis.

Prior studies have shown that heritability is enriched in regions surrounding genes with tissue-specific expression^30^. Thus, we first evaluated the contribution of all common variants to BP heritability in regions of expressed genes. To do so we defined the ‘gene region’ as a gene body spanning the transcription start to the stop sites ±50kb (**Figure 2A**). Next, for each of the four BP-relevant tissues we performed partitioning heritability analysis (S-LDSC)^29^ using all common sequence variants in gene regions of each tissue’s top 10,000 genes (ranked by median gene expression from GTEx^31^) and BP GWAS summary statistics from the UK Biobank^32^. These and the following analyses attempt to explain only that fraction of the heritability measured by genetic variants. For these analyses, we used a standard analysis where BP was adjusted for age and sex as covariates^33^. First, across all four tissues we explain ∼65% heritability which is highly significant. Second, each of the four tissues showed highly significant and ∼50% contribution to heritability (**Figure 2B**), suggesting significant pleiotropy.

**Figure 2:**
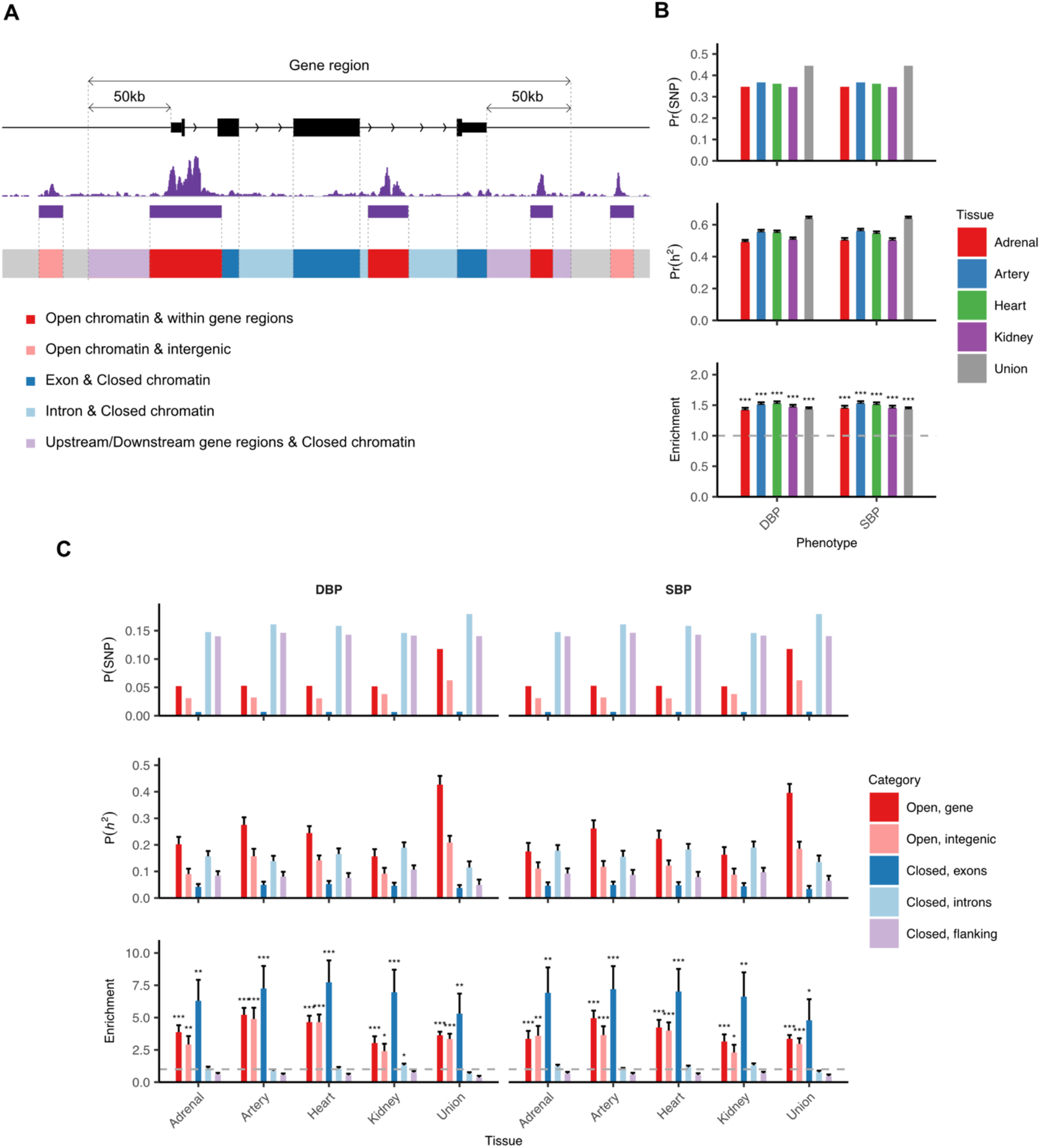
Variants in open chromatin regions are the major contributors to blood pressure heritability. **(A)** Schematic of how the genome is partitioned into five different non-overlapping regions based on gene annotation and chromatin accessibility. **(B)** Bar plots show the proportion of SNPs (top), the proportion of SNP heritability (middle), and their enrichment scores (bottom) for genes expressed in each of four tissues and their union. **(C)** For each tissue, the three statistics shown in (B) are further divided into the five exclusive genomic categories as described in (A).

We thus asked whether these contributions by variants in different functional categories in the gene regions are different. To understand these functional differences, we partitioned gene regions into five mutually exclusive and exhaustive subregions based on gene and CRE content, for each tissue (**Methods**). We evaluated heritability simultaneously for all subregions in one S-LDSC model so that we could directly compare their individual contributions in the context of other annotations. These analyses showed that CRE variants explain most of the heritability, particularly for the artery (27.5% DBP, 26.2% SBP) and heart (24.5% DBP, 22.3% SBP) while exonic variants showed a much smaller fraction (4.1 - 5.2% DBP, 4.4 - 4.8% SBP across tissues). In contrast, exonic variants showed the most significant enrichment (6.3 - 7.7 fold with *P* =1.2 × 10^−3^ - 8.2 × 10^−5^ for DBP; 6.6 - 7.2 fold with *P* = 2.7 × 10^−3^ - 5.2 × 10^−4^ for SBP across tissues) owing to its smaller target size (0.7% of all SNPs) (**Figure 2C**). Finally, we note that a significant portion of the heritability is explained by CREs outside expressed genes as well (9.0 - 15.7% DBP; 8.7 – 11.7% SBP, across tissues). This effect is likely from distal enhancers or CREs residing within genes expressed in other tissues. Although greater heritability is enriched in artery and heart CREs than in other tissues, we observed a high degree of explained heritability and enrichment in all subregions regardless of the tissue considered. This implies that variants in diverse functional elements cause inter-individual BP variation.

We hypothesize that numerous widely expressed genes and CREs active in multiple tissues, hereafter referred to as ubiquitous, may lead to significant BP heritability (**Figure S6A, B**). To test this contention, we first quantified the pattern of sharing of gene expression vis-à-vis their CREs across the four BP-relevant tissues we studied. The overlap of expressed genes across tissues demonstrates that more than 60% of them are expressed in all four tissues, while <5% are uniquely expressed (**Figure S6 C**). In contrast, 17% of CREs are ubiquitously open while 15-20% are open in each tissue only (**Figure S6D**). We further investigated this surprising relationship between gene expression and CREs across tissues by classifying CRE overlap between tissues vis-à-vis gene expression overlap. In general, the proportions of CREs with tissue overlaps do not vary much by gene expression overlaps (**Figure 3**). This is because two major CRE classes emerge; those that are ubiquitously open and those that are open only in the tissue in which its target gene is expressed. Consequently, we sought to quantify the contributions of CREs and genes to BP heritability, as a function of their ubiquitous vs. non-ubiquitous feature. For each tissue, we classified CREs into six different groups based on their gene expression and CRE activity patterns across tissues (**Figure 4A**) and performed partitioned heritability analysis using all six groups in one model. The most significant and greatest BP heritability arises from ubiquitously active CREs in ubiquitously expressed genes. This is the primary reason why we can explain significant heritability on an individual tissue basis (11.1 – 12.0% for DBP, 11.1 – 11.9% for SBP). On the other hand, BP heritability explained by tissue-restricted CREs in tissue-restricted genes is much more variable across tissues. Here, artery-(6.0% DBP, 6.5% SBP) and heart-restricted (5.2% DBP, 3.9% SBP) CREs explain much greater BP heritability than those from the adrenal gland (1.5% DBP, 2.1% SBP) and kidney (2.4% DBP, 2.2% SBP). Finally, we explained the greatest BP heritability enrichment in artery-restricted CREs for artery-restricted genes (6.7-fold with *P* =5.7 × 10^−5^ for DBP, 7.3-fold with *P* =2.8 × 10^−5^ for SBP). Similar to the result in **Figure 2C**, artery- and heart-restricted CREs not associated with “expressed genes” explain an even greater proportion of heritability (12.3% and 8.7% for DBP and SBP for artery and 8.7% and 8.5% for DBP and SBP for heart). Thus, distal enhancers play an important role in BP regulation, at least for the artery and heart.

**Figure 3:**
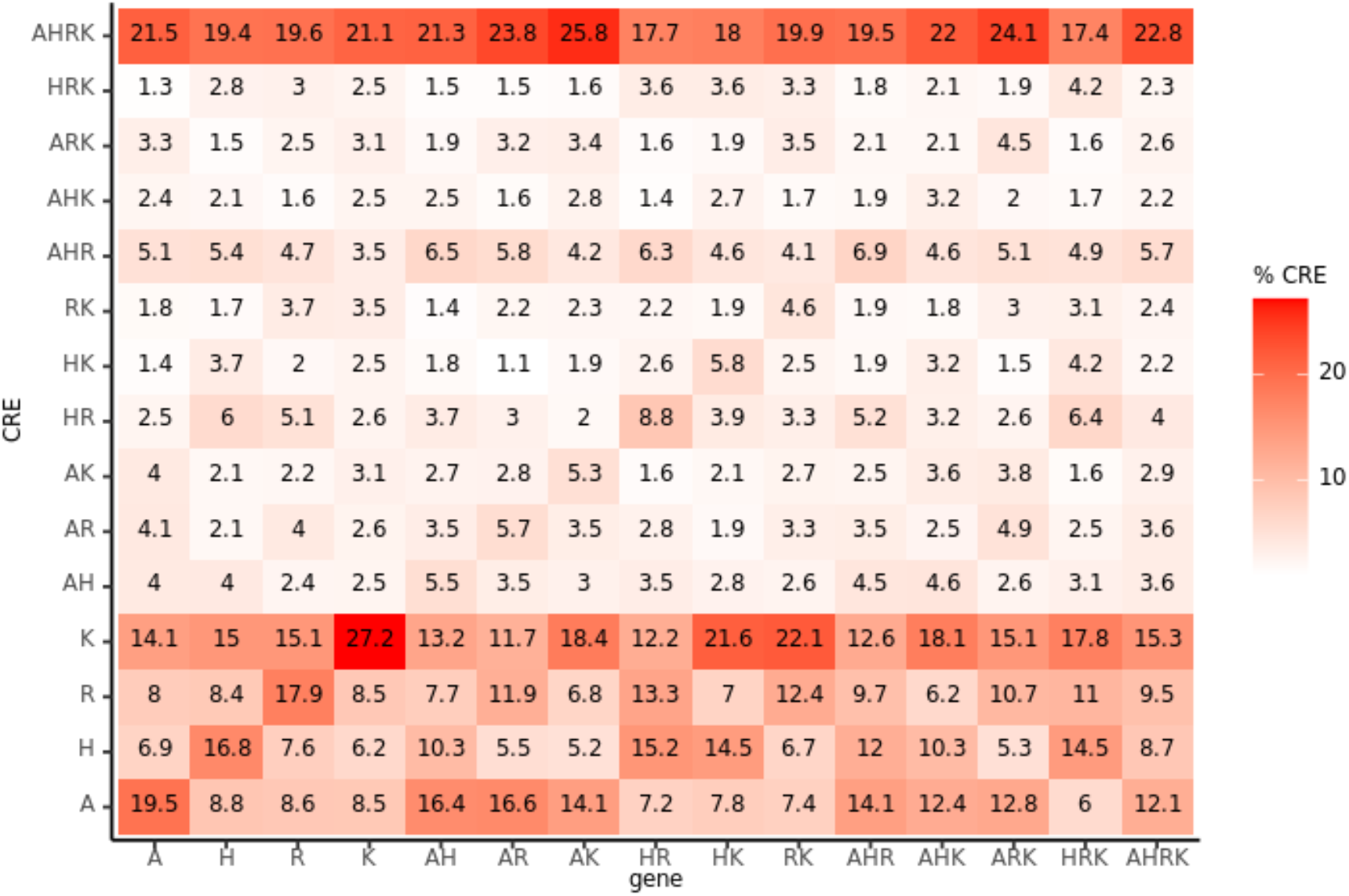
Open chromatin patterns are independent of gene expression patterns. For each of the gene expression patterns across four tissues (X axis), CREs are independently stratified by their open chromatin patterns across these same four tissues. A: Adrenal, R: Artery, H: Heart, and K: Kidney.

**Figure 4:**
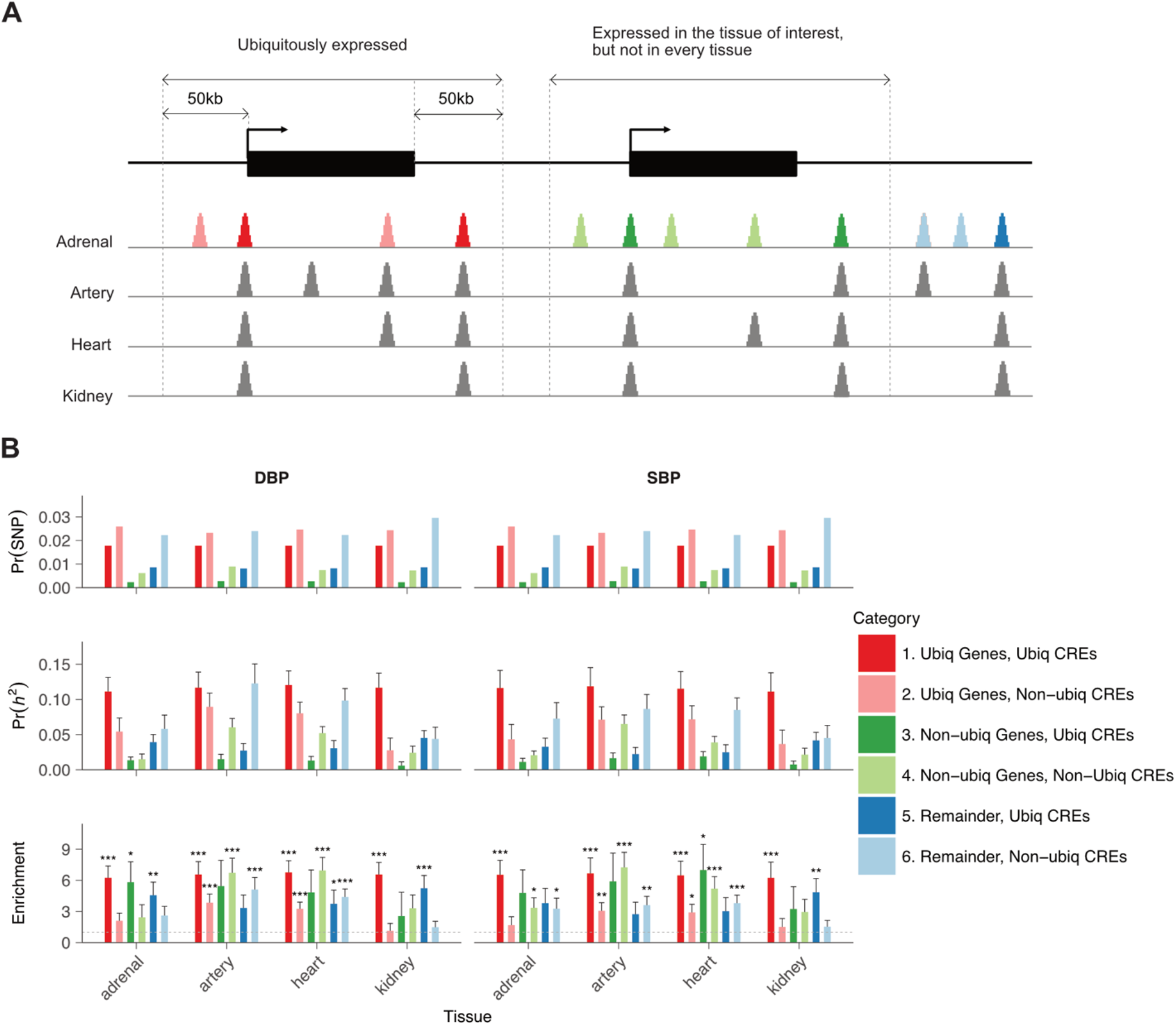
Non-ubiquitous CREs show the greatest differential enrichment scores for SBP and DBP. **(A)** Using adrenal gland as an example, we show how CREs active in adrenal gland can be classified into six different classes based on their activities in other tissues, genomic location and gene expression. **(B)** For each tissue, bar plots compare the proportion of SNPs (top), the proportion of heritability (middle), and the enrichment scores (bottom) across these six classes.

To disentangle the specific contribution of specific genomic regions across these BP-relevant tissues, we re-performed partitioning heritability analysis with different grouping of genomic regions. We considered all tissues together to define six mutually exclusive classes based on their overlap; four tissue-specific groups (adrenal gland, artery, heart, kidney), one common group (common), and one intermediate group (mixed). We then estimated the heritability explained by each of these groups in a single S-LDSC model. Consistent with our prior analysis, ubiquitously expressed genes explained the greatest BP heritability (41.4% DBP, 41.1% SBP) (**Figure 5A**); furthermore, CREs active in multiple tissues (common and mixed) explained the greatest heritability (15.0% DBP, 14.5% SBP for common and 19.0% DBP, 19.7% SBP for mixed) followed by artery (13.2% DBP, 10.1% SBP), heart (7.8% DBP, 4.9% SBP), and adrenal gland (5.5% DBP, 6.1% SBP) tissues (**Figure 5B**).

**Figure 5:**
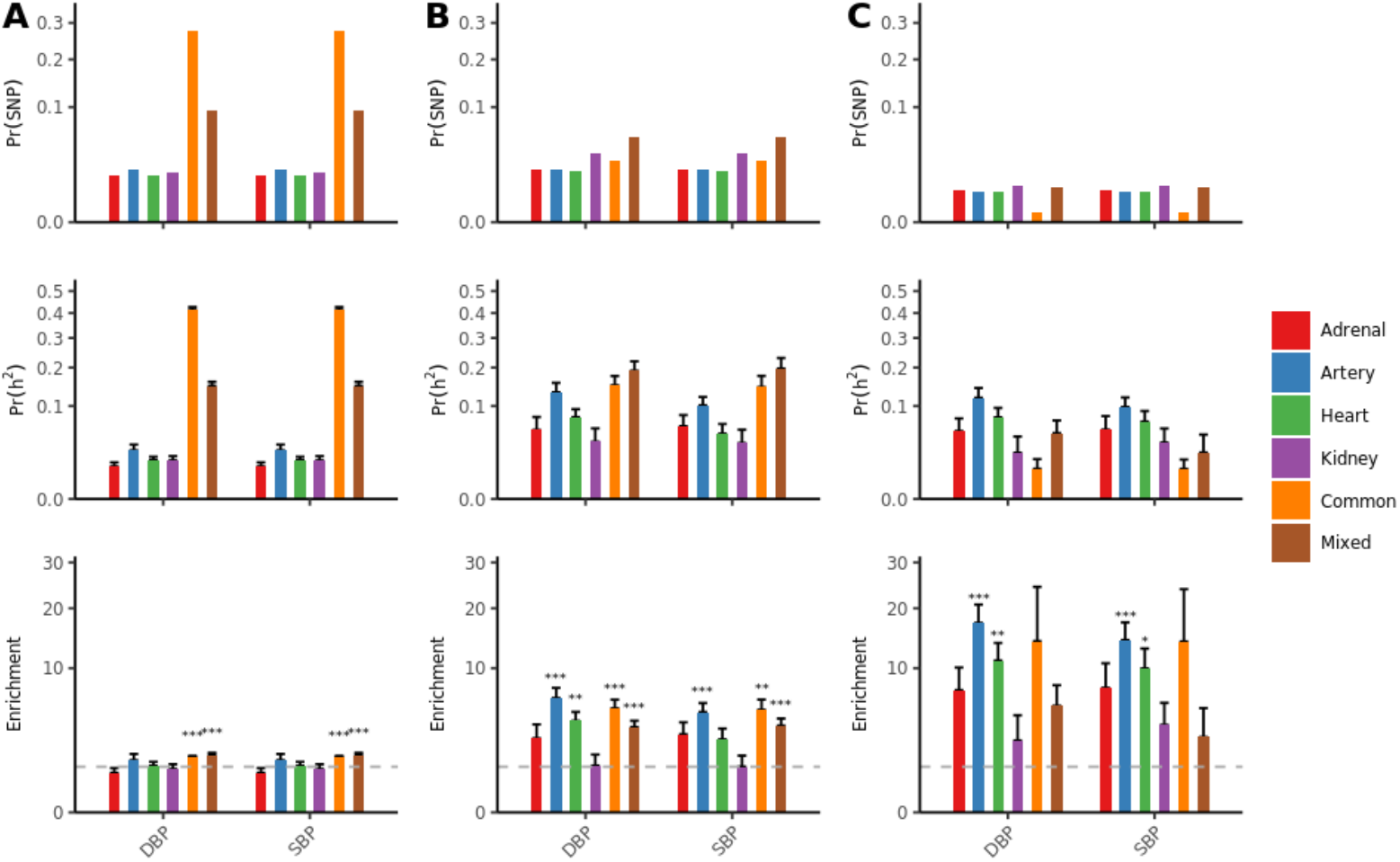
Predicted regulatory variants are largely tissue-specific and differentially affect SBP and DBP. Three different groups of variants are considered: (A) all variants in regions around expressed genes; (B) all variants in CREs; and (C) regulatory variants predicted by deltaSVM. In each group, variants were further divided into six different classes based on overlaps across the four tissues (see Figure 3). Variants in the “Common category” affect all four tissues while variants in the “Mixed category” affect more than one but not all tissues. For each variant group, bar plots show the proportion of SNPs (top), the proportion of heritability explained (middle), and the enrichment score (bottom) across the six variant classes.

Since our epigenomic maps could identify the tissue in which a CRE was active, regulatory variants within these CREs would indicate a tissue-specific genetic effect despite the CRE being ubiquitously open. We used the deltaSVM method with models trained on tissue-specific chromatin accessibility (**Figure S7**)^11^, to computationally predict the regulatory effect of variants in CREs (**Methods**). Strikingly, we discovered that most of these variants were predicted to have an impact on CREs in one tissue only. Specifically, among the variants in CREs in the four tissues, 15.9% and 30.2% are in common and mixed groups, respectively, while 1.8% and 22.3% of deltaSVM positive variants among all deltaSVM positive variants are in common and mixed groups. Moreover, these predicted regulatory variants have different levels of impact on BP phenotypes. Consistent with tissue-specific CREs, artery- (11.8% DBP, 9.8% SBP) and heart-specific (7.7% DBP, 6.9% SBP) regulatory variants explained the greatest BP heritability. Artery-specific regulatory variants achieved the most significant and the greatest enrichment (17.2 fold with *P* =5.3 × 10^−6^ for DBP; 14.2 fold with *P* =2.5 × 10^−5^ for SBP) (**Figure 5C**). Thus, even if regulatory elements are active in multiple tissues, variants in these regions are likely to exert their effect on phenotypes through a specific tissue. Collectively, we explain 33.4% and 29.5% of DBP and SBP heritability with these predicted regulatory variants in the four tissues.

As indicated earlier, regulatory variants that alter chromatin accessibility do so by disrupting TF binding. Therefore, we evaluated which TF motifs are enriched for the high-scoring 11-mers from the trained gkm-SVM models^26^ (**Methods**). This analysis revealed 137 TFs significantly enriched in at least one tissue (enrichment score ≥ 4) as well as expressed in that tissue (**Figure 6**). Interestingly, even though we only required TFs to be expressed in the tissue in which the corresponding motifs are enriched, most of the identified TFs (124 out of 137 TFs) did show expression in multiple tissues with differential enrichment across tissues. Moreover, while some TFs are uniquely enriched in one tissue, in general, most show enrichment in two or more tissues, suggesting that TFs work in combination with other TFs and that their activities are highly cell type- and tissue-dependent^34,35^. The most enriched TFs for artery are NFI (NFIA, NFIX, NFIB), AP1 (JUND, FOS), CEBP, and MEF2 (MEF2A/B/C/D) factors. In the heart, nuclear receptors (PPARA, NR2F2, NR4A1), NFI, and MEF2 are the most enriched, but MEIS, GATA and SOX family factors also demonstrated significant enrichment. We also found that motifs enriched for regions overlapping predicted regulatory variants (**Figure 6**) are largely concordant with the motifs enriched for the predictive 11-mers from the SVM models, suggesting that dysregulation of and by these TFs in their respective tissue of action affects BP regulation.

**Figure 6:**
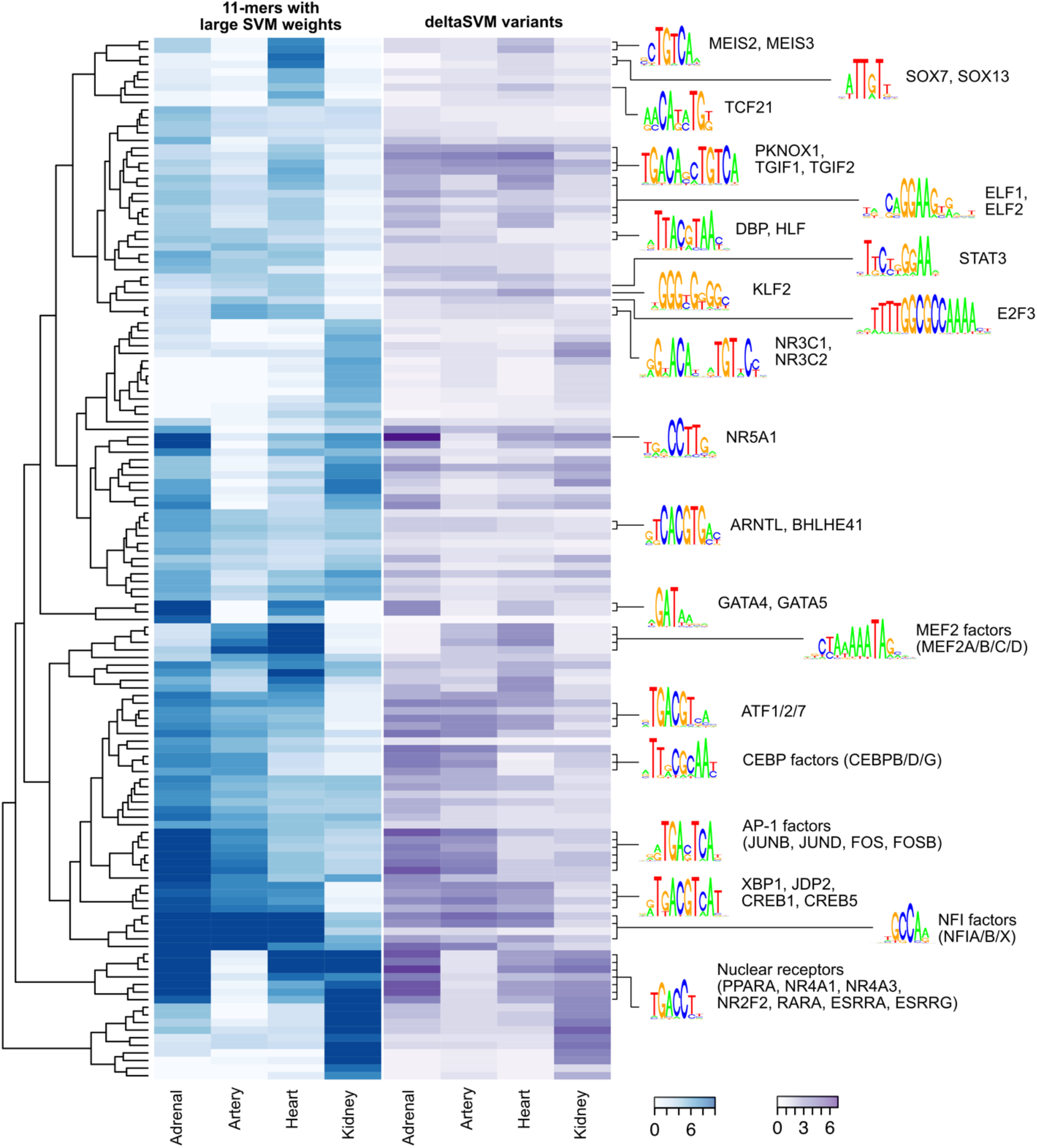
Specific transcription factors are differentially enriched for predicted regulatory variants. Heatmaps showing enrichment of transcription factor (TF) motifs across four tissues in 11-mers with large SVM weights from the trained SVM models (left) and in regions overlapping predicted regulatory variants (right). Motifs were clustered based on their enrichment pattern across tissues in the left. The motifs in both heatmaps are ordered the same. Representative TF motifs enriched in artery and heart are highlighted.

## Discussion

This study confirms that BP heritability is significantly controlled by genes expressed in artery, heart, kidney, and adrenal tissues and that inter-individual BP variation is largely controlled by CREs in distal enhancers. Although many of these genes are pleiotropic, surprisingly, CRE-mediated genetic effects from regulatory variants show far greater tissue-specificity. In other words, the pleiotropic nature of gene expression is largely caused by combinations of tissue-specific CREs. Therefore, regulatory sequence variants are more tissue-specific than the expression patterns that their genes indicate. Although exonic variants are many fewer than non-coding variants, we discovered that BP heritability shows the greatest enrichment in them (**Figure 2C**). This suggests that common exonic variants do affect BP phenotypes and provide an important route for identifying specific BP genes. Since the vast majority of these common exonic variants are not pathogenic, we suspect that they exert their effect on gene expression through differential codon usage of synonymous variants and missense variants affecting gene expression and function. We note that many of these exonic variants may also be tissue-restricted because only expressed genes with variants could exert their function in the tissue. Although not highly enriched, we also observe non-trivial BP heritability explained by intronic variants (but not in open chromatin regions). This suggests the emerging functional importance of deep intronic splicing variants on gene expression as well.

Of the four tissues studied, arterial and cardiac tissues have a greater role in BP heritability (**Figure 5**). The strong genetic effect of artery-specific regulatory variants on BP regulation is consistent with our previous finding that BP GWAS variants are most significantly colocalized with expression-modulating variants in artery tissues^36^. We have long known of two major ways to control BP, through cardiac output or through vascular resistance^37^. The arterial effect implicates genetic variation in vascular resistance is the greater contributor to inter-individual BP differences, and the specific mechanisms of this finding will require further investigation.

High-quality epigenomic maps are essential components of our approach. We showed that the quality of CRE maps is directly correlated with estimated BP heritability and that optimization of chromatin data processing further improves the yield of genetic analyses. Our approach is general and can be extended to other phenotypes. It is also straightforward to extend this approach to single-cell-based genomic annotations. More generally, our approach can now be used to answer broader physiological questions about BP, namely, how is the genetic architecture of systolic (SBP) and diastolic blood pressure (DBP) different, and what is the genetic basis of their correlation?

The enrichment analyses of TF motifs in the predictive sequence features and putative regulatory variants provide us an unique opportunity for mechanistic understanding of the genetic architecture of BP regulation. Our finding suggests that disruption of tissue-specific TF binding sites enriched for CREs are the major source of inter-individual variation of blood pressure. We anticipate that this is a common feature of many complex traits and diseases.

A major unanswered question in the genetics of multifactorial traits is the mechanism by which environmental factors modulate these traits. One possibility is that environmental factors affect these same tissues, dysregulate the same TFs and their cognate CREs to thereby dysregulate the same genes. In other words, genetic variation in CREs to alter specific target genes could be phenocopied by altering the TFs that bind these CREs. This hypothesis is biologically plausible because it proposes that BP variation arises from the same genes whose biology can be changed in specific tissues by both genetic and environmental factors but through different biological molecules: CREs versus TFs. The results from this study are critical to test this hypothesis. For example, the discovered functional architecture could be studied for its reaction to environmental exposures and to answer whether disease results through the *same biochemical and physiological pathways* irrespective of whether the perturbations are genetic or environmental.

## METHODS

### RNA-seq data processing (GTEx V8)

To identify expressed genes, we used the pre-processed GTEx RNA-seq datasets (http://gtexportal.org)^31^. Specifically, we obtained median gene-level TPMs by tissue (GTEx_Analysis_2017-06-05_v8_RNASeQCv1.1.9_gene_median_tpm.gct.gz) and labeled the top 10,000 genes ranked by the median TPM as “expressed.” For the kidney, we merged the cortex and medulla kidney data sets. We defined gene regions as 50kb extensions from both sides of gene bodies (transcription start and stop sites). The hg19 genome build and the gene annotation file from GENCODE v19 database were used^38^. The full list of genes is provided in **Data S1**. The genomic coordinates for these annotations are available in **Data S2**.

### ATAC-seq for kidney tissues

Four human kidney tissues were obtained as snap-frozen specimens from The National Disease Research Interchange (NDRI) as well as from the University of Michigan and Gift of Life Michigan. We employed two different protocols as a part of our experiment optimization. The ATAC-seq libraries for the first two samples (R1 and R2) were generated using the protocol adapted from Corces et. al.^39^ Briefly, tissue samples were finely chopped on ice and homogenized in an Accumax solution (Sigma-Aldrich) using a Dounce homogenizer. The samples were then passed through a 100um cell strainer followed by a 40μm cell strainer. The cells were pelleted in a cold centrifuge at 2,000 rpm and washed with cold PBS twice. We resuspended the pellet in 125μL of ATAC-Resuspension Buffer (RSB) containing 0.1% NP-40, 0.1% Tween-20 and 0.01% Digitonin, and incubated on ice for 3 minutes. We washed the lysis with 5ml ATAC wash buffer (10mM Tris-HCL, 10mM NaCl, 3mM MgCl2, 1% BSA, 0.1% Tween-20) and counted nuclei using trypan blue (1:9). We kept 50K nuclei from each sample, and centrifuged at 500 RCF (10 minutes, 4°C). Pellets were resuspended in 50ul of transposition mixture, containing 2.5uL transposase in 1x TD buffer (20034198, Illumina) with 0.001% Digitonin and 0.01% Tween 20, and incubated for 30 minutes on a thermomixer at 1000RPM and 37°C. DNA was extracted using MinElute Reaction Cleanup Kit (Qiagen).

The remaining two samples were processed using an improved protocol that includes sorting as follows. Tissues were finely chopped, washed with ice-cold PBS (w/o Ca2+/Mg2+) and homogenized in Nuclei EZ lysis buffer (NUC-101, Sigma Aldrich) as recommended in presence of 0.25U/μl RNAse inhibitor. Lysis was promoted by gentle mixing of homogenate using bore tips, followed by incubation for 5 minutes on ice and passing of homogenate through 70μm cell strainer. Nuclei was then pelleted by centrifugation of the filtered homogenate at 500g for 5 minutes at 4 degrees. Nuclei were washed twice with 500μl Nuclei Wash and Resuspension Buffer (1x PBS, 1% BSA, 0.25μl RNAse) for 5 minutes on ice (first wash without resuspending nuclei, second wash with resuspended nuclei). After each wash, nuclei were pelleted by centrifugation at 500g for 5 minutes at 4 degrees. Pelleted nuclei were further resuspended in 100-500μl Nuclei Wash and Resuspension Buffer with DAPI (10μg/ml) and nuclei were sorted by FACS to eliminate cellular debris as well as doublets. Sorted nuclei were then pelleted at 500g for 5 minutes at 4 degrees and 500μl ATAC wash buffer (10mM Tris-HCL, 10mM NaCl, 3mM MgCl2, 1% BSA, 0.1% Tween-20) was gently added and incubated for 5 minutes on ice without resuspending nuclei. Washed nuclei were then pelleted by centrifugation at 500g for 5 minutes at 4 degrees and nuclei was gently resuspended in 500μl ice-cold ATAC wash buffer and incubated for 5 minutes. Nuclei pellet was obtained by centrifugation at 500g for 5 minutes at 4 degrees and resuspended in 100μl ice-cold 1x TD buffer (20034198, Illumina). About 10K nuclei was used for transposition reaction at 37 degrees for 30 minutes in a thermomixer. DNA was extracted using MinElute Reaction Cleanup Kit (28206, Qiagen). Sequencing was carried out at the Genome Technology Center, New York University and 50 million 150bp paired-end reads per sample were obtained for each sample.

### DNase-seq & ATAC-seq data processing

For CRE map construction, we primarily used chromatin accessibility data from public databases (**Table S1**). Specifically, we collected DNase-seq and ATAC-seq samples from ENCODE^27^ for the adrenal gland, heart (left ventricle), and tibial artery. We also generated ATAC-seq libraries for adult kidney tissues, for which no public data existed (see above).

We uniformly processed DNase-seq and ATAC-seq data using our previously established pipeline with a minor modification^26^. Starting with properly paired and non-duplicated read pairs, we extracted the cut-sites from each read and used them as input to the MACS2 peak caller^40^ with the no model option (--nomodel). We found the optimal extension and shift base-pairs to be 100bp (--extsize 100) and -50bp (--shift -50; lagging strand), respectively. Also, we used --keep-dup option and default q-value cut-off (*q* <0.01). For ATAC-seq samples, we additionally adjusted the cut-sites by +4bp for forward-strand reads and -5bp for reverse-strand reads to take into account the 9bp insertion by the Tn5 transposase^41^.

### Partitioning heritability using S-LDSC

We obtained GWAS summary statistics for systolic (S) and diastolic (D) blood pressure (BP) phenotypes from the UK Biobank^32^ as processed by the Ben Neale laboratory (http://www.nealelab.is/uk-biobank/). To estimate SNP-heritability 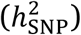 from GWAS summary statistics, we used a stratified version of LD-score regression (S-LDSC) to estimate the proportion of SNP-heritability of SNPs in a category as previously described^29,42^. Briefly, we primarily used enrichment scores to estimate the relative contribution of SNPs in specific annotations (e.g., tissue-specific CREs) to heritability. The enrichment score is calculated as proportional SNP heritability contributed to a SNP set (Pr(*h*^2^_SNP_)), divided by the proportion of SNPs in that SNP set (Pr(SNP)). We used z-scores from normalized S-LDSC coefficients to compare samples, which correspond to the statistical significance of the per-SNP contribution to the heritability as previously reported^29^.

We defined genomic annotations of CREs as ±1,000bp extended regions from peak summits to capture putatively functional variants in flanking regions associated with CREs^26^. To control signals confounded by other functional regions, such as protein-coding and evolutionary-conserved regions, we used the 97 baseline LD model as recommended^43^. We used data from European ancestry subjects and corresponding allele frequencies from the 1000 Genomes Phase 3 data as a reference panel for LD score calculation. We obtained the baseline model and reference panel data sets from https://data.broadinstitute.org/alkesgroup/LDSCORE/. For a fair comparison among multiple annotations of interest, we conducted S-LDSC analyses of all annotation sets in one model along with the baseline model.

### gkmQC

We assessed the quality of open chromatin peaks using gkmQC^28^. Specifically, we split the open chromatin peaks into subsets, each comprising 5,000 peaks sorted by decreasing signal intensity scores from the peak caller. We then trained the gkm-SVM model for each peak subset and used the “predictability” of peaks from models as a quality metric of peak subsets. Model performance was measured by the area under the ROC curves with five-fold cross-validation (i.e., AUC scores of the peak subsets). To visualize overall quality of a sample, we plot AUC scores as a function of ranks of peak subsets (i.e., gkmQC curve). We limit the analysis to the top 100,000 peaks (=20 subsets of 5,000 peaks each) to generate a gkmQC curve. We use the 10^th^ AUC score of peak subsets as an overall sample quality and determine a sample to be high-quality if the 10^th^ AUC is greater than 0.8.

### gkm-SVM

We built gkm-SVM models following our previously established pipeline with minor modifications^25,26^. For each high-quality sample as determined by gkmQC, we defined the positive training set as follows: starting from the top 100,000 open chromatin regions (ranked by their MACS2 p-values obtained from our optimized pipeline described above), we removed from the training set peaks with >1% of N-bases, >70% of repeats, and commonly open regions (defined as regions active in at least 30% of samples across all ENCODE data sets), as previously described^11,26^. We further restricted open chromatin regions to overlapping H3K27ac peaks from the same tissue (**Table S2**). As a negative training set, we used an equal number of random genomic regions, matched for length, GC content and repeat fraction of the positive set. To prevent potential biases caused by variable sequence length, we used 600bp fixed-length regions as a training set by extending ±300bp from peak summits. We used LS-GKM^25^ software for training with l=11, k=7, d=3, and t=4 (weighted-gkm kernel). For each sample, we averaged ten different models with different random samplings of negative training sets. After training, we further combined the models from different samples (i.e., biological replicates) to generate one model per tissue.

### deltaSVM

For each tissue, we calculated deltaSVM scores for all common variants (∼10M with minor allele frequency > 1% in the EUR superset) from the 1000 Genome Project (v3)^44^, using the combined model and identified variants overlapping open chromatin regions in the corresponding tissue. We used ±10 bp regions centered on those variants for scoring. We then determined variants in the top 15th percentile of deltaSVM scores as potentially functional regulatory variants. This threshold was chosen based on our previous analyses of allele-biased chromatin accessibility in heart tissues^26^.

### TF motif enrichment analysis

We identified TF motifs enriched for high-scoring 11-mers (i.e., the top fifth percentile) from the trained gkm-SVM models, as previously described^26^ with a minor modification. Briefly, for each of the manually curated TF motifs from the CIS-BP database^45^, we identified all 11-mers that significantly matched a given motif using FIMO with default parameters^46^, and calculated the enrichment of the matched 11-mers in the top fifth percentile ranked by their SVM weights. We determined TF motifs as significant if their multiple testing corrected binomial *P* ≤ 0.01 and enrichment score was ≥ 4. To further reduce false positives, we only considered TFs expressed in the tissue, as previously defined (i.e., the top 10,000 genes ranked by median gene expression from GTEx). Similar to enrichment analysis of the high-scoring 11-mers, we also identified TF motifs enriched for regions surrounding predicted regulatory variants. We used ±10 bp regions centered at each variant to scan the TF motifs using FIMO with default parameters. We tested both reference and alternative alleles and determined ‘a hit’ if either of them significantly matched a motif. We calculated the expected hit frequency using all common variants and the observed hit frequency using the predicted regulatory variants only. The enrichment scores were then calculated as (observed hit frequency) / (expected hit frequency).

### Data access

All sequencing reads generated in this study have been submitted to the NCBI Gene Expression Omnibus (GEO; http://www.ncbi.nlm.nih.gov/geo) under accession number GEOXXXXXX.

## Acknowledgements

We thank Dr. Luciano Marteletto for his advice and improved protocol for the ATAC-seq experiment. This study has benefited from useful comments from Drs. Xiaofeng Zhu, Alanna Morrison, and Charles Gu. This research was supported by the computational resources of the high-performance computing core at NYU and National Institutes of Health grants HL086694, HL141980 and HL128782 to A.C..

## Author contributions

D.L. and A.C. conceived and designed the study; S.K.G. collected the adult kidney tissues; O.Y., H.B., and P.M. performed ATAC-seq experiments; D.L. and S.K.H. conducted all computational analyses; and, D.L., S.K.H., and A.C. wrote the manuscript. All authors were involved in manuscript revision.

