## Supplemental Material for "Tissue-specific and tissue-agnostic effects of genome sequence variation modulating blood pressure"

[#]Corresponding authors:

### This document includes:

Supplemental Figures S1 to S8

Supplemental Tables S1 to S3

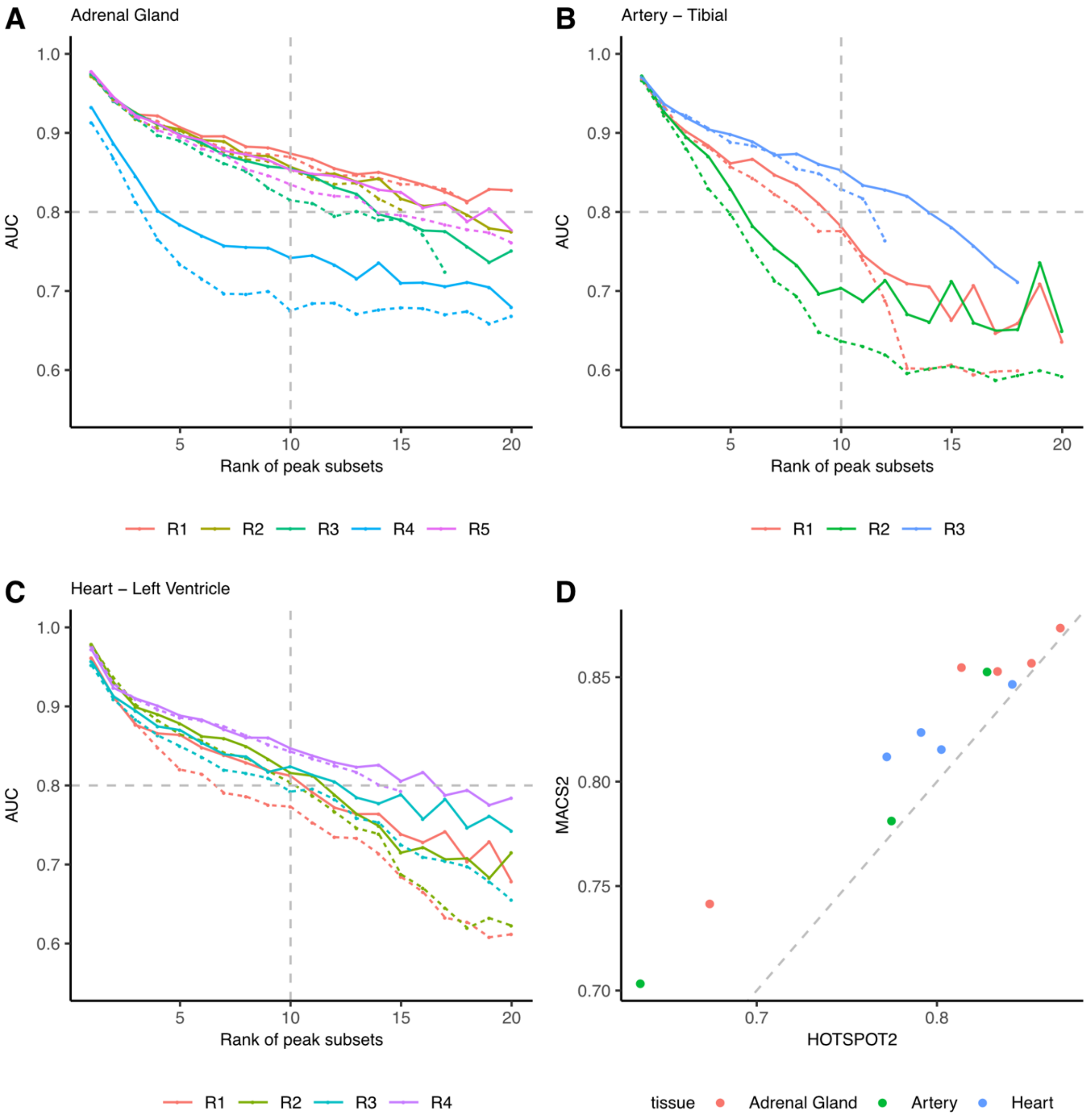


**Figure S1: MACS2 peak calling quality is superior to HOTSPOT2.** We compared the quality of accessible chromatin peaks between two different peak calling algorithms (MACS2 and HOTSPOT2) for DNase-seq using gkmQC. For each sample, we recalled peaks from the processed bam files using MACS2 while HOTSPOT2 peaks were obtained from the ENCODE portal. The gkmQC plots, which compares peak predictability (AUCs) across peak subsets stratified by their peak signal strengths, are shown for **(A)** Adrenal Gland, **(B)** Artery - Tibial, and **(C)** Heart - Left Ventricle. **(D)** Per-sample AUC of the $10^{th}$ peak subset is compared between the two peak callers. MACS2 peak sets achieve consistently better gkmQC scores across nearly all samples regardless of tissue.

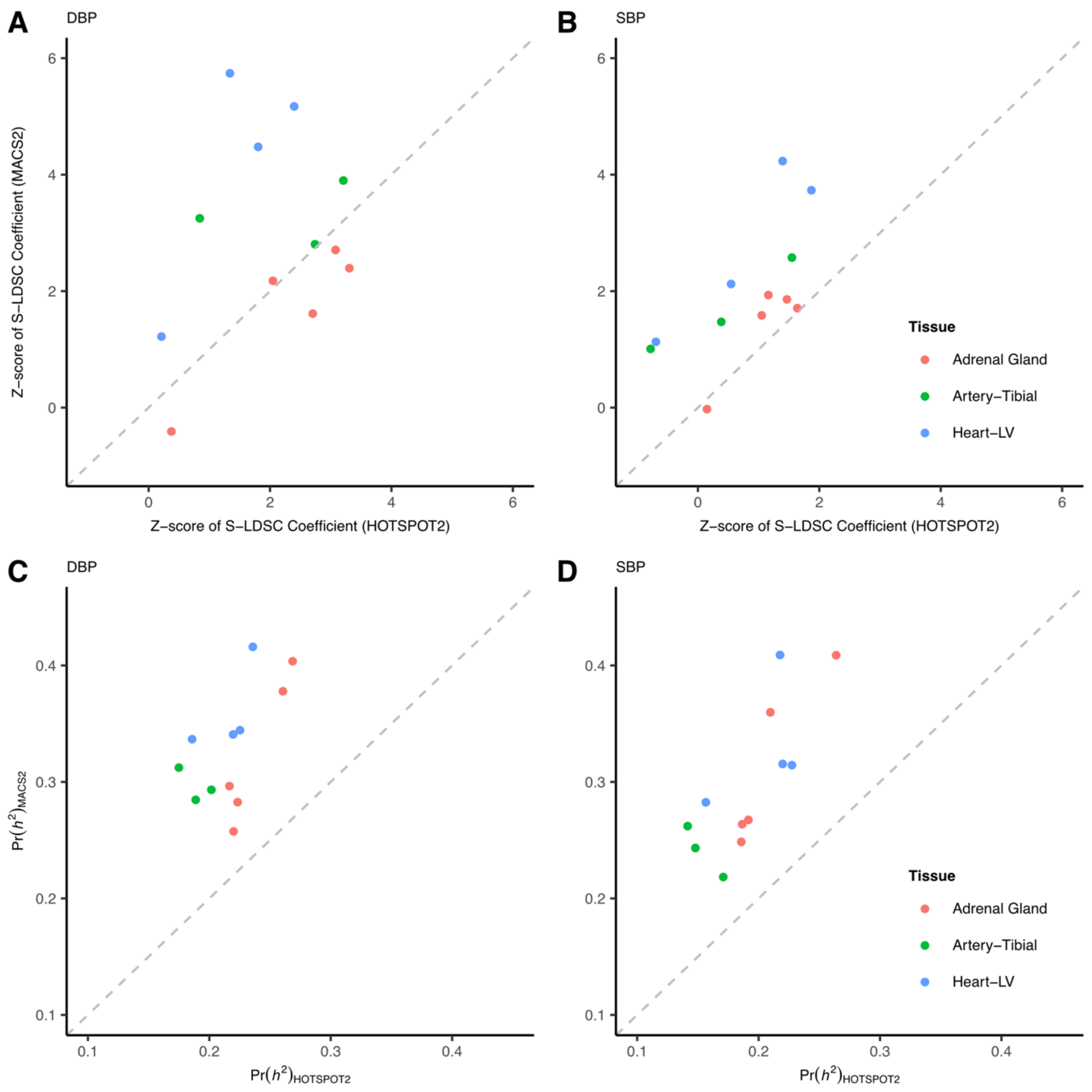


**Figure S2: Variants within MACS2 peaks explain greater blood pressure heritability than HOTSPOT2.** For each DNase-seq sample, Z-scores of S-LDSC coefficients, corresponding to the statistical significance of per-SNP heritability, are compared between the two peak sets for **(A)** Diastolic (DBP) and **(B)** Systolic (SBP) blood pressure. **(C, D)** Similarly, the proportion of SNP-heritability ($h^2$) is compared between the two peak sets for SBP and DBP. Consistent with **Figure S1**, variants in MACS2 peaks make greater contributions to BP heritability than variants in HOTSPOT2 peaks.

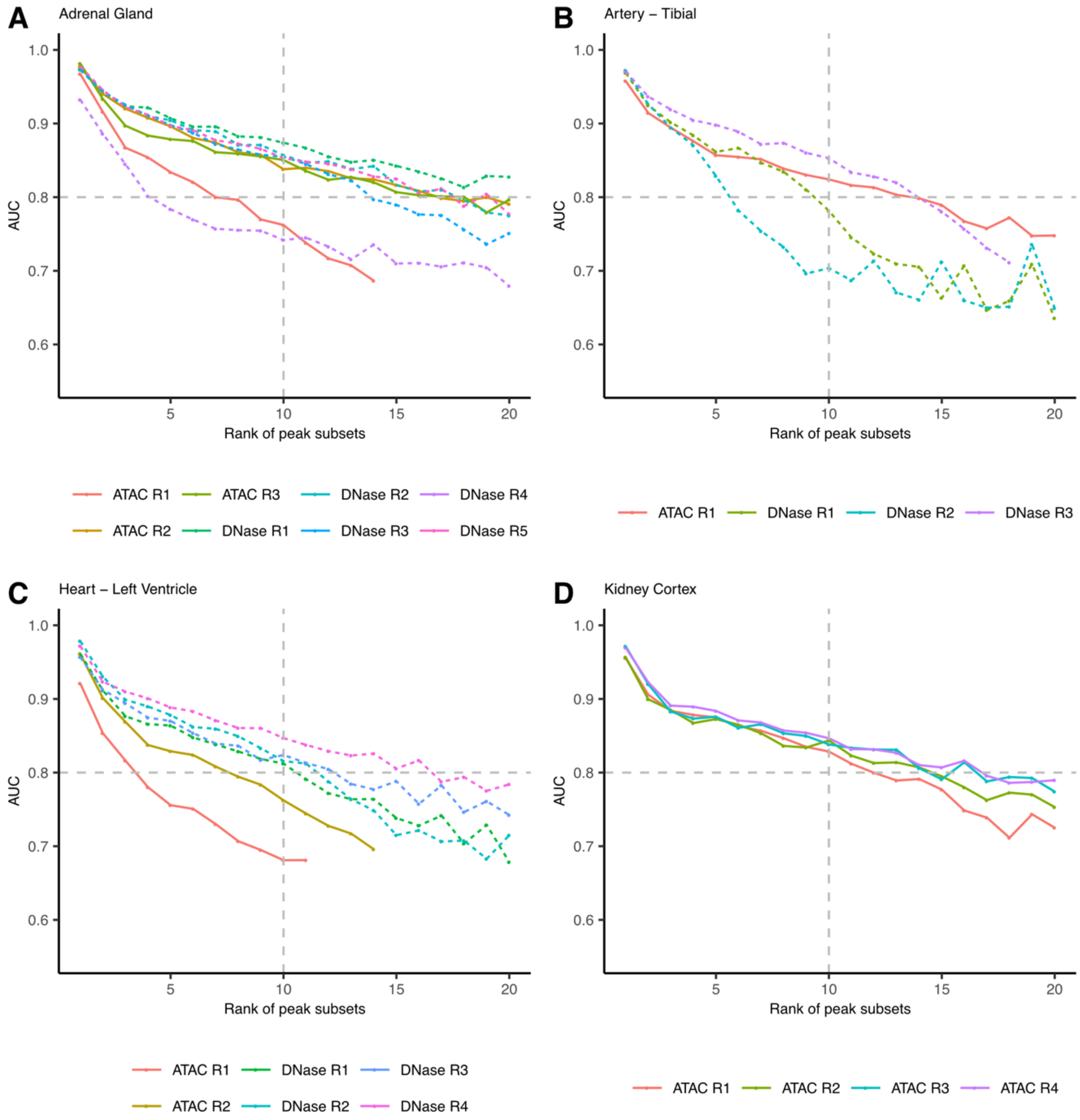


**Figure S3: gkmQC analyses of chromatin accessibility data from four tissues relevant to BP regulation.** To maximize coverage of chromatin accessibility peaks in the genome, we also analyzed ATAC-seq data sets from the ENCODE project for **(A)** Adrenal Gland, **(B)** Artery-Tibial, and **(C)** Heart-LV. **(D)** We generated four ATAC-seq data sets for adult kidney tissues. All four data sets were assessed by gkmQC. ATAC-seq samples are displayed as solid lines. For comparison, DNase-seq samples are shown as dotted lines. In total, six adrenal gland samples, two tibial artery samples, four heart left ventricle samples, and four kidney cortex samples passed our gkmQC criteria (AUC>0.8 at the 10$^{th}$ peak subset).

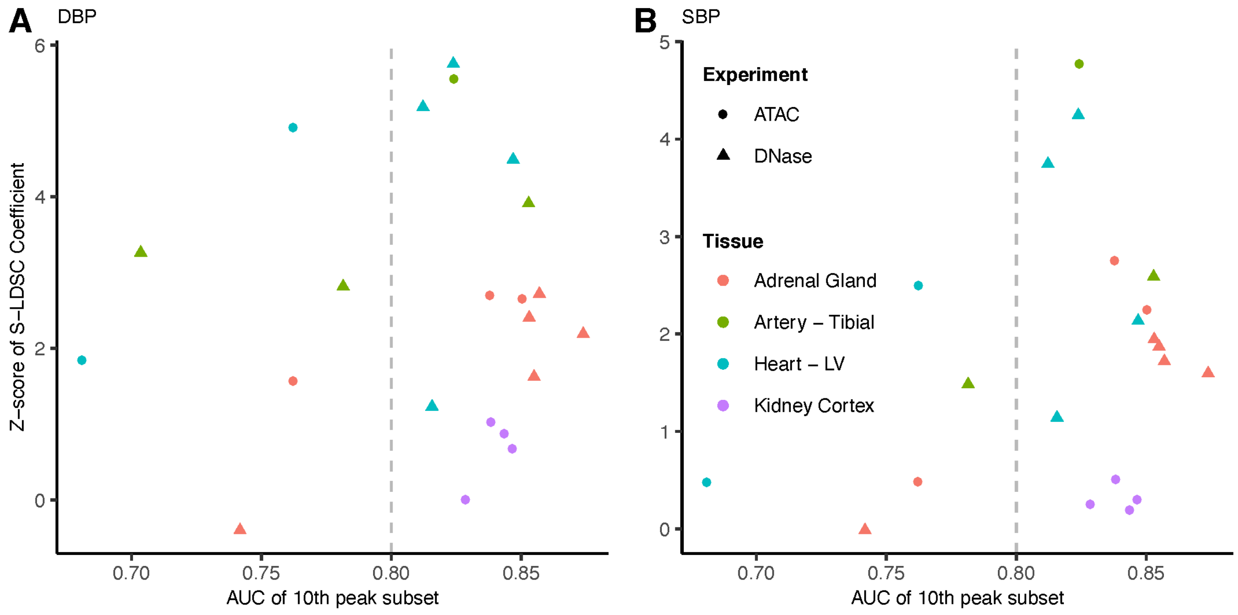


**Figure S4: gkmQC scores correlate with blood pressure heritability**. For each of the samples from the four BP-relevant tissues, the AUC of the 10$^{th}$ peak subset (X-axis) is compared to the Z-score of an S-LDSC coefficient (Y-axis) for **(A)** Diastolic (DBP) and **(B)** Systolic (SBP) blood pressure.

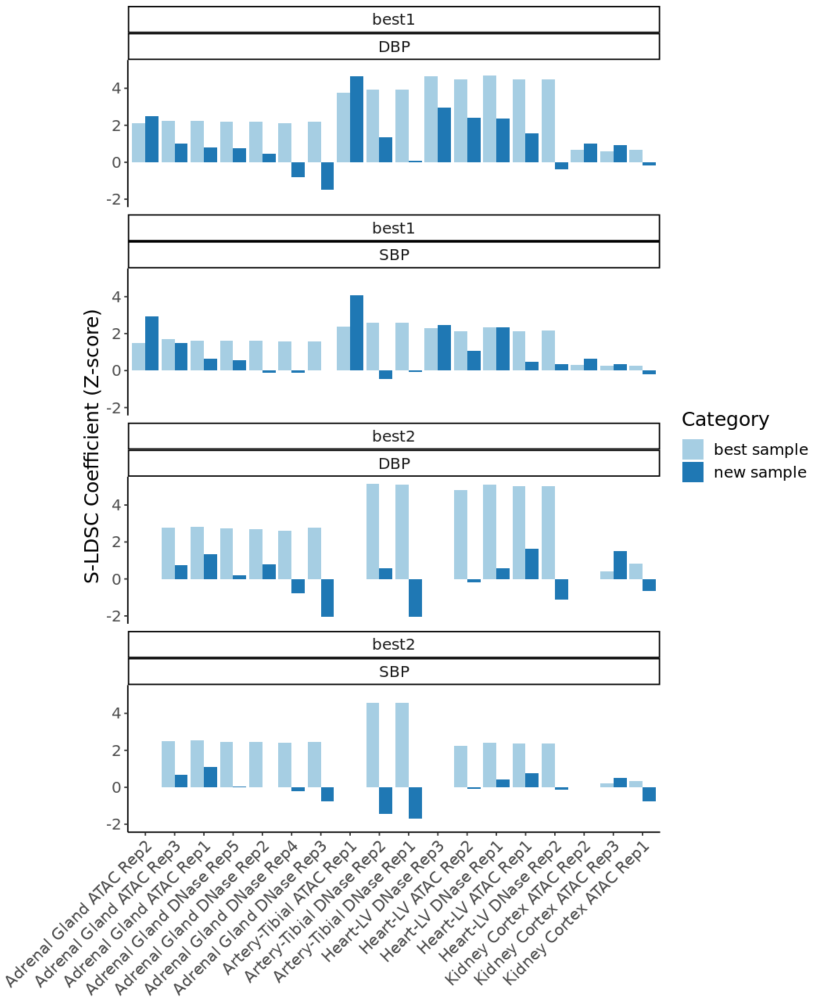


**Figure S5: Different types of chromatin accessibility data improve chromatin accessibility map quality. (1st and 2nd rows):** For each tissue, we performed partitioned heritability analysis of DBP and SBP using chromatin accessibility peaks uniquely detected in a sample after controlling for peaks observed in the best sample, as determined by gkmQC. Light blue bars are Z-scores of S-LDSC coefficients for peaks from the best sample, and the adjacent dark blue bars are those for peaks uniquely detected in a given sample. The best samples are Adrenal Gland DNase Rep1, Artery DNase Rep3, Heart-LV DNase Rep4, and Kidney Cortex ATAC Rep 4, all of which are UW DNase-seq datasets. Except for the kidney tissues, where we only had ATAC-seq data, we found that at least one sample exhibited higher Z-scores than the best sample and that these samples were either ATAC-seq or DUKE DNase-seq. **(3rd and 4th rows)** The same analysis was repeated using the combined set of peaks from the two best samples as a new control. Z-scores of S-LDSC coefficients for the combined sets were more significant than peaks from the single-best samples. Peaks uniquely observed in the remaining samples capture nominal heritability (Z-score <1.5) after controlling for peaks in the best two samples combined.

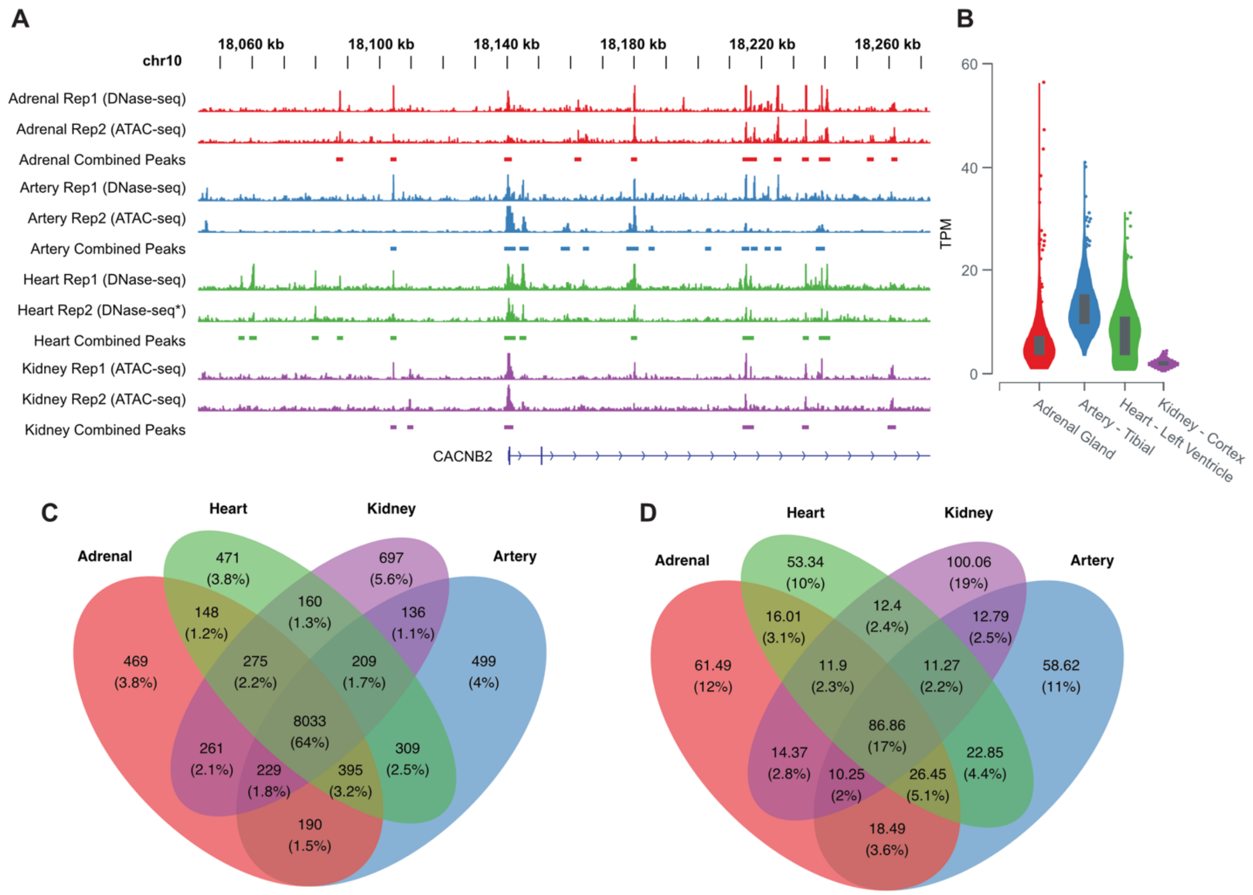


**Figure S6: Chromatin accessibility is more tissue-specific than gene expression.** **(A)** Chromatin accessibility patterns across the four tissues demonstrate that many open chromatin regions are tissue-specific; the example shows *CACNB2* which is significantly associated with blood pressure. **(B)** *CACNB2* is expressed in all four tissues; violin plots show gene expression distribution (TPM) of *CACNB2* across individuals from GTEx V8. **(C)** Venn diagram shows overlap of gene expression across the four tissues together with the number of shared genes and their proportions. **(D)** Venn diagram shows overlap of open chromatin regions across the four tissues together with the total bases of overlap (MB) and their genomic proportion.

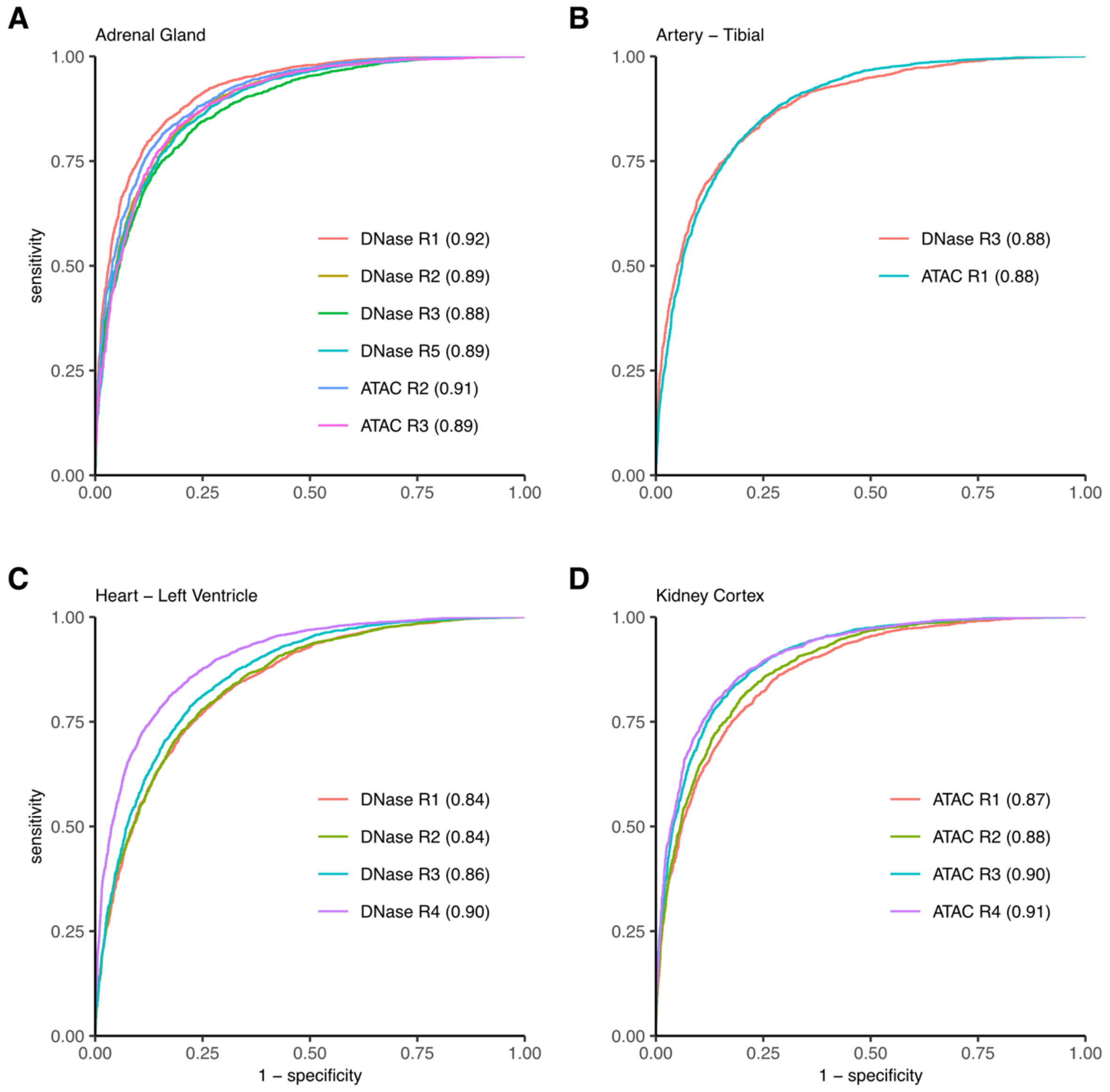


**Figure S7: gkm-SVM accurately predicts tissue-restricted open chromatin regions.** We followed the previously established framework to build a tissue-specific gkm-SVM model for each of the high-quality samples and used peaks not common to all samples. ROC curve analyses for 20% of reserved peaks are shown for **(A)** Adrenal gland, **(B)** Artery-Tibial, **(C)** Heart-LV, and **(D)** Kidney Cortex. The area under the ROC curves (AUC) are shown in parentheses.

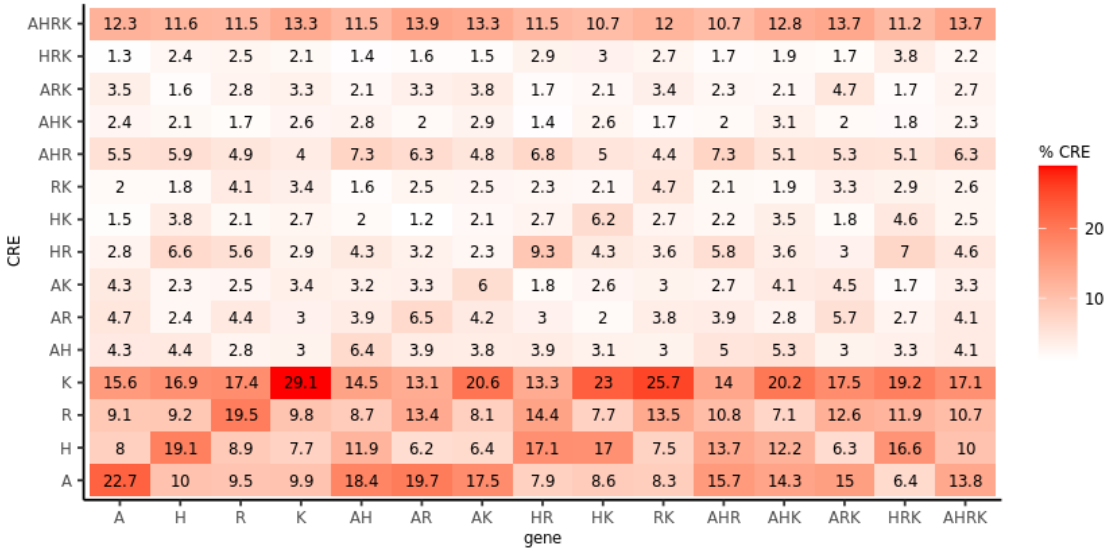


**Figure S8: Analysis of Figure 3 after removing promoter proximal regions (defined as ±1000bp from the nearest transcription start site).** For each of the gene expression overlap groups (X axis), CREs are independently stratified by their chromatin accessibility across the four tissues. A: Adrenal, R: Artery, H: Heart, and K: Kidney.

| Tissue | Replicate Name | ENCODE bam file accession | ENCODE Hotspot2 file accession | Experiment | gkmQC | Final |
|---|---|---|---|---|---|---|
| Adrenal Gland | DNase Rep1 | ENCFF429WIJ | ENCFF721ZHK | DNase-seq (UW) | Yes | Yes |
| | DNase Rep2 | ENCFF647DDQ | ENCFF335CNN | DNase-seq (UW) | Yes | |
| | DNase Rep3 | ENCFF830ETB | ENCFF654WWP | DNase-seq (UW) | Yes | |
| | DNase Rep4 | ENCFF961UNE | ENCFF977OWF | DNase-seq (UW) | | |
| | DNase Rep5 | ENCFF994CAF | ENCFF350XGZ | DNase-seq (UW) | Yes | |
| | ATAC Rep1 | ENCFF374OSU | . | ATAC-seq | | |
| | ATAC Rep2 | ENCFF711SUI | . | ATAC-seq | Yes | Yes |
| | ATAC Rep3 | ENCFF436NOT | . | ATAC-seq | Yes | |
| Artery-Tibial | DNase Rep1 | ENCFF091MJM | ENCFF601CWS | DNase-seq (UW) | | |
| | DNase Rep2 | ENCFF656IUA | ENCFF048ZGK | DNase-seq (UW) | | |
| | DNase Rep3 | ENCFF817DBS | ENCFF620ZPK | DNase-seq (UW) | Yes | Yes |
| | ATAC Rep1 | ENCFF168OTV | . | ATAC-seq | Yes | Yes |
| Heart-LV | DNase Rep1 | ENCFF021QCV | ENCFF225UJM | DNase-seq (Duke) | Yes | |
| | DNase Rep2 | ENCFF037AJZ | ENCFF146VYU | DNase-seq (UW) | Yes | |
| | DNase Rep3 | ENCFF442IPX | ENCFF892RWC | DNase-seq (Duke) | Yes | Yes |
| | DNase Rep4 | ENCFF837QAV | ENCFF850WOE | DNase-seq (UW) | Yes | Yes |
| | ATAC Rep1 | ENCFF906UQF | . | ATAC-seq | | |
| | ATAC Rep2 | ENCFF310VHO | . | ATAC-seq | | |
| Kidney Cortex | ATAC Rep1 | . | . | ATAC-seq | Yes | |
| | ATAC Rep2 | . | . | ATAC-seq | Yes | Yes |
| | ATAC Rep3 | . | . | ATAC-seq | Yes | |
| | ATAC Rep4 | . | . | ATAC-seq | Yes | Yes |

**Table S1: List of chromatin accessibility data sets used in this study.** We focused analyses on four BP-relevant tissues. Chromatin accessibility data sets for Adrenal Gland, Tibial Artery, and Heart Left Ventricles were obtained from the ENCODE project while adult kidney ATAC-seq data were generated in our laboratory.

| Tissue | ENCODE bed narrowPeak accession | Number of peaks |
|---|---|---|
| **Adrenal Gland** | ENCFF115DDQ | 34,734 |
| | ENCFF193BKF | 21,409 |
| | ENCFF511YRX | 18,727 |
| | ENCFF852GEC | 38,480 |
| | ENCFF985YQH | 28,228 |
| **Artery-Tibial** | ENCFF066YNA | 41,219 |
| | ENCFF377PYQ | 46,918 |
| **Heart-LV** | ENCFF170EQJ | 49,199 |
| | ENCFF543XLN | 45,526 |
| | ENCFF094TOI | 37,209 |
| | ENCFF258ZRC | 47,560 |
| **Kidney Cortex** | ENCFF584JKX | 33,134 |
| | ENCFF512JNT | 37,336 |

**Table S2: List of H3K27ac ChIP-seq data sets used in this study.** We focused analyses on four BP-relevant tissues; pseudo-replicated peaks were obtained from the ENCODE project.

| Tissue | # CREs | Average length (bp) | Genome coverage (%) |
|---|---|---|---|
| **Adrenal Gland** | 126,996 | 2380 | 9.64 |
| **Artery-Tibial** | 109,351 | 2438 | 8.50 |
| **Heart-LV** | 117,835 | 2359 | 8.86 |
| **Kidney Cortex** | 160,797 | 2428 | 12.45 |
| **Combined** | **245,008** | **2698** | **21.07** |

**Table S3.** Characteristics of the high-quality chromatin accessibility maps we generated for four BP-relevant tissues.
